# Hypercholesterolaemia promotes epigenetic memory in hematopoietic stem cells that persists after lipid lowering and causes systemic immunometabolic dysfunction mediated by macrophage metabolic reprogramming

**DOI:** 10.1101/2025.09.15.676283

**Authors:** Gareth S.D. Purvis, Jiahao Jiang, Thomas K. Hiron, Christina Simoglou Karali, Matthew Baxter, Rong Li, Rohit Vijjhalwar, Marina Diotallevi, Sally A. V. Draycott, Surawee Chuaiphichai, Gillian Douglas, Eileen McNeill, Adam J. Mead, David R. Greaves, Chris A. O’Callaghan, Keith M. Channon

## Abstract

In patients with atherosclerotic cardiovascular disease (ASCVD) lowering LDL cholesterol (LDL-C) reduces the risk of adverse cardiovascular events. However, these patients remain at continued risk of adverse cardiovascular events despite achieving optimal LDL-C lowering. We hypothesised that this ‘residual risk’ is mediated by epigenetic programming of haematopoietic stem cells (HSC) which persists after cholesterol levels are lowered and results in sustained effects on innate immune cell metabolism, that is not resolved by cholesterol lowering. We found that exposure to high cholesterol (HC) *in vivo* induced long-term metabolic changes in macrophages differentiated *ex vivo* from bone marrow, leading to a pro-inflammatory profile; these changes persisted *in vivo* despite cholesterol lowering. HSC from HC mice had altered chromatin accessibility that persisted after cholesterol lowering and was also present in bone marrow monocytes and tissue resident macrophages. HC provoked RUNX1-dependent downregulation of stearoyl-CoA desaturase (SCD) which reduced mono-unsaturated fatty acid (MUFA) availability for OxPhos in murine and human monocytes and macrophages. Supplementation with MUFA restored OxPhos capacity and promoted a shift towards a less pro-inflammatory macrophage phenotype in HC-trained BMDM. Bone marrow chimera and lineage tracking studies revealed that prior exposure of HSC to HC conferred adverse systemic metabolic effects on normocholesterolemic mice, with increased adipose tissue mass and increased migration of macrophages derived from HC-exposed HSC into adipose tissue, resulting in increased adipose tissue inflammation and systemic glucose intolerance. These findings indicate that HC results in long-lasting immuno-epigenetic memory in HSC which is refractory to lipid lowering, and provide strong evidence that exposure to HC can have prolonged consequences that require new therapeutic approaches, beyond cholesterol lowering.

## Introduction

High cholesterol is a major risk factor for atherosclerotic cardiovascular disease (ASCVD)^1^. Interventions to lower LDL-cholesterol (LDL-C) dramatically reduce the risk of major adverse cardiovascular events (MACE) such as myocardial infarction. People with, or at high risk of, ASCVD now routinely receive treatment to reduce LDL-C, most frequently with statins ^2^. Treatment with high dose statins and newer LDL-lowering drugs such as PCSK9 inhibitors can produce major reductions in LDL-C (<1 mM or 40 mg/dl) ^3,4^ However, there is limited incremental benefit from further lowering of LDL-C with a ‘residual risk’ remaining even after aggressive reductions in LDL-C^4^. Therefore, identifying other modifiable determinants of residual cardiovascular risk is a crucial step to achieve future therapeutic benefit.

Metanalysis of 31,245 ASCVD patients treated with statins in three major multinational trials (PROMINENT, REDUCE-IT, and STRENGTH), demonstrated that inflammation is a major contributor to residual risk, as assessed by high-sensitivity C-reactive protein ^5^. This suggests that there is a substantial component of residual risk, mediated by inflammation, that is not mitigated by reducing cholesterol levels ^6^, but could be targetable if the molecular mechanisms mediating this residual risk could be identified.

Immunological memory is central to the adaptive immune response. However, innate immune cells may also retain a memory of a previous exposure to an activating stimulus, via epigenetic reprogramming which influences disease progression ^7^. A hallmark of innate trained immune response is bioenergetic re-programming towards Warburg-type metabolism. Innate trained immunity can be induced by systemic cardiometabolic abnormalities such as obesity ^8^ or hyperglycaemia ^9^ in mice, and can promote disease progression ^10^. Hyperglycaemia ^9^ or an intermittent high fat diet ^11^ can induce trained immunity in HSC leading to increased plaque formation ^12^. However, the role of innate trained immunity in the residual risk after cholesterol-lowering remains unexplored, and the impact of cholesterol-lowering on other tissues, beyond vessels, that are important in cardiometabolic health is also unknown. In particular, adipose tissue inflammation is a key driver of the adverse systemic consequences of obesity ^13^, and a CV risk factor independent of LDL-C ^8^.

Accordingly, we sought to identify consequences of prior exposure to HC that persist in HSC and myeloid cells after cholesterol-lowering, and to determine their effects on systemic metabolic health and adipose tissue function. This revealed HC-induced trained immunity in HSC and myeloid cells, and provides a rational basis for developing new approaches to target residual risk in ASCVD.

## Methods

### Animal studies

All animal studies were conducted with ethical approval from the Local Ethical Review Committee at the University of Oxford and in accordance with the UK Home Office regulations (Guidance on the Operation of Animals, Scientific Procedures Act, 1986). Mice were housed in ventilated cages with a 12-h light/dark cycle and controlled temperature (20–22□°C) and fed normal chow and water *ad libitum* unless otherwise stated. Mice had received no prior procedures (including acting as breeding stock) prior to use in experiments for this manuscript.

#### Model of inducible hypercholesterolaemia

Male *w*ild-type or hCD68-GFP mice were injected with a single dose of 3 ×10^11^ adeno-associated virus 8 (AAV8) encoding a mutated form of the proprotein convertase subtilisin/kexin type 9 (mPCSK9). Following injection mice were fed a high-fat diet (HFD) (D12079B) from Research diets, USA. Control mice received a single dose of 3×10^11^ AAV8 control particles and maintained in chow diet ^14^.

#### High cholesterol-lowering intervention

Hypercholesterolaemia was induced in male wild-type CD57BL/6 mice using a single intraperitoneal injection of AAV8-mPCSK9 and feeding with a HFD. After 12 weeks a cohort of mice would undergo high cholesterol-lowering intervention (HCLI) and be reverted onto a chow diet and given a mono-clonal anti-body against PCSK9 (Evolocumab; 10 mg/kg *s.c.* weekly), to lower lipid level to baseline.

#### Bone marrow transplant

Hypercholesterolaemia was induced in male hCD68-GFP mice using a single intraperitoneal injection of AAV8-mPCSK9 and feeding with a HFD. After 12 weeks bone marrow is harvested. Control mice received control AAV8. 7-week-old wild-type mice were irradiated with a non-lethal dose of 10G, administrated in 2 doses separated by 4 hours. Irradiated WT mice then received 5×10^6^ sex-matched donor bone marrow cells. Two weeks after BMT, chimerism efficiency was tested; only mice with above 90% GFP+ blood neutrophils were used for subsequent studies. In some studies BMT-mice were fed a high-fat high-sugar (HFHS) diet (D12331i) from Research diets, USA to induce diet induced obesity (DIO) ^15^.

#### Oral glucose tolerance test

The assay was performed as previously described ^16^, briefly, mice were fasted for 6 hours prior to the start of the assay, at which point baseline measurements of blood glucose levels were taken from the tail vein. Mice were then given an oral dose of glucose (2 mg/kg), and blood glucose measured at 15 min intervals for 120 mins.

#### Zymosan induced peritonitis

Hypercholesterolaemia was induced in hCD68-GFP mice using a single intraperitoneal injection of AAV8-mPCSK9 and feeding with a HFD. After 12 weeks mice were challenged with zymosan (Zymosan A; 100□μg; i.p.) ^17^. PECs were harvested in 7 ml of ice-cold PEC harvest buffer (PBS, 5-mM EDTA) 2 after zymosan challenge.

### Cell Culture of primary macrophages

#### Murine Bone Marrow-Derived Macrophages (BMDMs)

Bone marrow-derived macrophages were generated as previously described. Briefly, fresh bone marrow cells from tibiae and femurs of male mice were isolated and cultured in Dulbecco’s Modified Eagle’s medium (DMEM) supplemented with 10% heat-inactivated fetal bovine serum (FBS), 10% L929 cell-conditioned media as a source of macrophage colony-stimulating factor, and 50□U/ml penicillin/streptomycin for 7□days. Bone marrow cells were seeded into 8□ml of medium in 100□mm non-tissue culture treated Petri dishes (Thermo Fisher Scientific, Sterilin, UK). On day 5, an additional 5□ml of medium was added. Gentle scrapping was used to lift cells off the dish surface. BMDMs were then counted and resuspended in DMEM containing 2 % FBS and 1 % penicillin/streptomycin media at the desired cell concentration for experiments.

#### Resident Peritoneal Macrophages

The peritoneal cavity was lavaged with 7□ml ice-cold PBS supplemented with 5□mM EDTA. Cells were pelleted by centrifugation, resuspended in DMEM 10% FBS, and plated for 2–4□h to allow macrophages to attach to the plate. Non-adherent cells were washed off.

### Seahorse XFe96 analysis of mitochondrial function

On day 7 of culture, macrophages were plated at a density of 7.5 × 10^4^ cells per well into XFe96 microplates. Cells were left to attach for 2 hours before stimulation with LPS/IFNγ or IL-4 if required, for 18 hours. Extracellular flux analysis was then performed. One hour prior to the assay, cells were washed and the culture medium was replaced with Seahorse XF Base Medium (modified DMEM without Phenol Red, pH 7.4; Agilent), supplemented with glucose (10 mM, Sigma), glutamine (2 mM, Sigma), and sodium pyruvate (2 mM, Sigma), before being incubated at 37°C, at atmospheric CO2 levels. Oxygen consumption rate (OCR) and extracellular acidification rate (ECAR) were measured using the XFe96 analyser (Agilent). 6 base-line OCR measurements were taken followed by 3 measurements after sequential injection of the following compounds 2 μM, oligomycin (Sigma), 2 μM FCCP (Sigma), and combined 0.5 μM antimycin A (Sigma), and 0.5 μM rotenone (Sigma). Cells were measured in 4 replicate wells and all data normalized to cell number. For calculations the third reading in each case was used as the most stable point, except for FCCP where the highest reading was taken.

### Metabolomics

BMDM were grown in 9 cm non-tissue-treated plates, stimulated with 100 ng M(IL-4) for 18 hours before being pelleted and frozen on dry ice and stored at 80C. Metabolon measured metabolites using mass spectrometry, as outlined below. Samples were prepared using the automated MicroLab STAR system from Hamilton Company. Several recovery standards were added prior to the first step in the extraction process for QC purposes. To remove protein, dissociate small molecules bound to protein or trapped in the precipitated protein matrix, and to recover chemically diverse metabolites, proteins were precipitated with methanol under vigorous shaking for 2 min (Glen Mills GenoGrinder 2000) followed by centrifugation. The resulting extract was divided into five fractions: two for analysis by two separate reverse phase (RP)/UPLC-MS/MS methods with positive ion mode electrospray ionization (ESI), one for analysis by RP/UPLC-MS/MS with negative ion mode ESI, one for analysis by HILIC/UPLC-MS/MS with negative ion mode ESI, and one sample was reserved for backup. Samples were placed briefly on a TurboVap (Zymark) to remove the organic solvent. The sample extracts were stored overnight under nitrogen before preparation for analysis.

#### Ultrahigh Performance Liquid Chromatography-Tandem Mass Spectroscopy (UPLC-MS/MS)

All methods utilized a Waters ACQUITY ultra-performance liquid chromatography (UPLC) and a Thermo Scientific Q-Exactive high resolution/accurate mass spectrometer interfaced with a heated electrospray ionization (HESI-II) source and Orbitrap mass analyzer operated at 35,000 mass resolution. The sample extract was dried then reconstituted in solvents compatible with each of the four methods. Each reconstitution solvent contained a series of standards at fixed concentrations to ensure injection and chromatographic consistency. One aliquot was analysed using acidic positive ion conditions, chromatographically optimized for more hydrophilic compounds. In this method, the extract was gradient eluted from a C18 column (Waters UPLC BEH C18-2.1×100 mm, 1.7 mm) using water and methanol, containing 0.05% perfluoropentanoic acid (PFPA) and 0.1% formic acid Another aliquot was also analysed using acidic positive ion conditions, however it was chromatographically optimized for more hydrophobic compounds. In this method, the extract was gradient eluted from the same afore mentioned C18 column using methanol, acetonitrile, water, 0.05% PFPA and 0.01% formic acid and was operated at an overall higher organic content. Another aliquot was analysed using basic negative ion optimized conditions using a separate dedicated C18 column. The basic extracts were gradient eluted from the column using methanol and water, however with 6.5mM ammonium bicarbonate at pH 8. The fourth aliquot was analysed via negative ionisation following elution from a HILIC column (Waters UPLC BEH Amide 2.1×150 mm, 1.7 mm) using a gradient consisting of water and acetonitrile with 10mM ammonium formate, pH 10.8. The mass spectroscopy analysis alternated between MS and data-dependent MSn scans using dynamic exclusion. The scan range varied slighted between methods but covered 70-1000 m/z.

### ELISA

Supernatant from cell culture was diluted so cytokines were detectable with the range of the kit. Capture anti-body was bound to Nunc MaxiSorp ELISA plates, and incubated overnight at 4 C before being blocked (0.05 % BSA, PBS). After washing diluted supernatant and standards was loaded and incubated at room temperature with rocking for 2 hours. After washing, plates were incubated with detection antibody at room temperature with rocking for 2 hours. After washing, plates were incubated with HRP conjugates secondary antibody for 30 mins and washed. Plates were then incubated with 50 µl TMB substrate for 20 mins in the dark, before stop solution was added directly to the plate. Optical density of each well was immediately read, using a microplate reader set to 450 nm. If wavelength correction was available it was set to 540 nm.

### Stromal vascular fraction (SVF) isolation

Adipose tissue was harvested into PSB supplemented with 2 % FBS and minced into 1 mm pieces using a scalpel. Tissue was then incubated in digestion media (100 mM HEPES, 2 % BSA, 1.5 mg/mL collagenase II in Hank’s Balanced Salts) doe 40 mins at 37 C using a shaker at 150 rpm. Digested tissue was the filtered into a single cell suspension using a 100 µM cell strainer, and spun at 100 G for 8 mins. The adipose layer was removed by aspiration, and the remaining cells spun at 400 G for 5 mins. Red blood cells were removed by addition of ACK lysis buffer for 3 mins. At which stage cell were spun and counted for experimentation.

### Flow cytometry

Cells were washed in FACS buffer (0.05 % BSA, 2 mM EDTA in PBS pH 7.4), blocked using anti CD16/32 for 10 mins at 4°C, followed by antibody staining for the cell surface markers with appropriate isotype controls. Fluorescence-minus-one (FMO) and single staining controls were used for gating and compensation. Absolute cell counts were quantified using counting beads added to each sample (CountBright, Invitrogen). Data were acquired using a BD Fortessa X20 cytometer and Diva software (BD Biosciences) and then analysed using FlowJo (Tree Star Inc, USA) software.

### Microscopy

2×105 BMDM were seeded onto glass cover slips prior to polarisation (M(LPS/IFNγ; 100 ng/mL LPS + 20 ng/mL) or M(IL-4); IL-4 20 ng/mL) for 18 hours. Mitochondria were stained using MitoTracker, according to the manufacturer’s instructions.

### Bone marrow and hematopoietic stem cells quantification and isolation

Flow cytometry of bone marrow: Cryo-stored bone marrow was defrosted rapidly in DMEM (2 % FBS and DNase I) and centrifuged at 300 RCF for 20 mins 4°C. Cells were counted and 10^7^ cells were stained to quantify hematopoietic lineage and stem populations. Cells were Fc-blocked in anti-CD16/32 for 10 mins at 4°C, and stained with anti-mouse antibodies. 7-Amino-Actinomycin D (7-AAD; Sigma) was used for dead cells exclusion. Fluorescence-minus-one (FMO) controls, isotype control antibodies and negative populations were used as gate-setting controls. FACS analyses were performed on BD LSRII or BD Fortessa X20 and subsequent data analyses were performed with the FlowJo analysis software. For HSC isolation, cryo-stored bone marrow was defrosted rapidly in DMEM (2 % FBS and DNase I) and centrifuged at 300 RCF for 20 mins 4°C. 2.5×10^7^ cells were Fc-blocked in anti-CD16/32 for 10 mins at 4°C, and stained with anti-mouse antibodies, 4′,6-diamidino-2-phenylindole (DAPI; Invitrogen) was used for dead cells exclusion. Cell sorting was performed on BD FACS AriaIII sorter (BD Biosciences), with a mean cell sorting purity of 97.4□±□0.45 % (mean□±□s.e.m.).

### HSC ATAC-sequencing

Cryo-stored bone marrow was rapid defrosted in DMEM (2 % FBS and DNase I), single cell suspensions were then stained for FACS. 500 Lin-Sca+ckit+ HSC’s were then directly sorted into transposition lysis buffer (containing TD buffer, 1 % digitonin and 10 % Tween-20), Tn5 tagmentation enzyme was added immediately after sorting and samples were incubated at 37 C for 30 minutes. Resultant fragmented DNA was cleaned up and eluted using the Qiagen MinElute Reaction Cleanup Kit. Libraries were indexed and amplified using the NEB Next High-fidelity PCR kit using customised Nextera primer pairs (https://www.nature.com/articles/nmeth.2688). Indexed libraries were sequenced using 150 bp pair end read by NovoGene.

### Bone marrow monocytes isolation and ATAC Sequencing

Cryo-stored murine bone marrow was defrosted into DMEM (2 % FBS and DNase I). Monocytes were then immunomagnetically sorted using Monocyte Isolation Kit (BM), mouse, according to the manufacture’s instructions (Miltenyi Biotec; 130-100-629). Cell viability was assessed, and 75,000 bone marrow monocytes were used. Briefly, 75,000 viable cells were centrifuged (500 RCF) at 4°C for 5 min and resuspended in 50 ul cold ATAC-Resuspension Buffer (RSB) supplemented with (0.1% NP40; 0.1% Tween20; and 0.01% Digitonin) for 3 min and washed in 1 ml ATAC-RSB supplemented with 0.1 % Tween-20. Nuclei were then pelleted at 500 RCF for 10 mins at 4°C in a fixed-angle centrifuge. The pelleted nuclei were then resuspended in 50 ul of transposition buffer Transposition mix (2x TD buffer, transposase (100nM final), PBS, 1% digitonin, 10% Tween-20) and incubated at 37°C for 30 min. Resultant fragmented DNA was cleaned up and eluted using Zymo DNA Clean and Concentrator-5 Kit (D4014). Samples were amplified for 5 cycles using the NEB Next High-fidelity PCR kit. DNA was cleaned up and eluted using Zymo DNA Clean and Concentrator-5 Kit. Libraries were then quantified using the KAPA Library Quantification kit (KK4824) and pooled libraries were sequenced using NextSeq 150 bp pair end read by NovoGene.

### ATAC data analysis

For HSC: Raw ATAC sequencing reads were processed using CATCH-UP ^18^ to generate bam files mapped to the mm10 genome. Bigwig tracks where merged and LanceOtron ^19^ used to call peaks. Differential binding analysis was performed using DiffBind v3.14.0 and PCA plots were generated using deepTools v3.5.2.

For BM monocytes: Raw ATAC sequencing read were aligned to the mm10 genome using Bowtiw2 ^20^. A consensus peak set was called using MACS3 ^21^ combining all samples. DiffBind v3.14.0 was used to for differential accessibility analysis, and TOBIAS^22^ for motif foot printing analysis.

### RUNX1 Chip-Seq

ChIP-seq data for RUNX1 in human cord blood CD34+ cells and in THP-1 monocytes were retrieved from the NCBI Gene Expression Omnibus. The accession numbers used were GSE111917 and GSE254674, respectively. Chromatin accessibility was assessed in the SCD loci, and peaks called using MACS3. Signals from technical replicates were merged for visualisation purposes and normalized by sequencing depth.

### Single-cell multiome

Previously generated single-cell multiome data from human monocyte-derived macrophages (NCBI Sequence Read Archive accession number PRJNA1102756). In summary, CD14+ monocytes were isolated from healthy human donors and differentiated into macrophages using M-CSF over a period of 7 days. This was followed by a 48-hour treatment with 50 ug/mL of ox-LDL or a control buffer. For the differential gene expression analyses, we used sequencing-depth-normalised expression, quantified as count-per-10k-reads. Data was then projected in UMAP and clustered as previously described ^23^, expression of SCD and RUNX1 was overlaid on individual cells showing maximal expression plots.

### Statistical Analysis

Statistical analyses were performed using Graph Pad Prism 10 (Graph Pad Software). Experimental groups were compared using two-tailed Student’s unpaired *t-*tests. When more than two groups were compared a one-way analysis of variance (ANOVA) for multiple comparisons was conducted with Dunnet’s correction. Data are presented as mean□±□s.e.m with individual data points plotted demonstrating biological replicates. *P*□<□0.05 was considered to be significant. Seahorse data is plotted as sum of 4 technical replicates.

## Results

### High cholesterol induces sustained abnormalities in macrophage metabolism and pro-inflammatory polarisation

To investigate the sustained effects of hypercholesterolaemia on macrophage function, we induced hypercholesterolaemia in mice, exposing bone marrow hematopoietic stem cells (HSC) to high cholesterol (HC) levels *in vivo.* As myeloid cells have a relatively short life span, we hypothesised that any persisting effect of previously high cholesterol levels would be mediated through HSC (Figure 1A). Intraperitoneal injection of an adeno-associated virus encoding PCSK9 with a gain-of-function D377Y mutation (AAV8-mPCSK9) followed by high fat diet (HFD) feeding was used to generate hypercholesterolaemia (Figure 1B). Control mice injected with a AAV8-null and maintained on chow diet did not have elevated cholesterol levels (Figure 1B). To assess whether the effects of high cholesterol are ‘memorised’, bone marrow-derived macrophages (BMDM) were differentiated *ex vivo* from hypercholesterolaemic or control mice under standard conditions. BMDM generated from HC and control mice showed no difference in basal activation state (Figure C-F). However, when stimulated with LPS/IFNγ BMDM from HC mice showed a striking exaggeration of the pro-inflammatory response compared with BMDM from control mice (Figure 1C/D). Conversely, when stimulated with IL-4 to induce a pro-resolution phenotype, BMDM from HC mice displayed a diminished response (Figure 1E/F).

**Figure 1:**
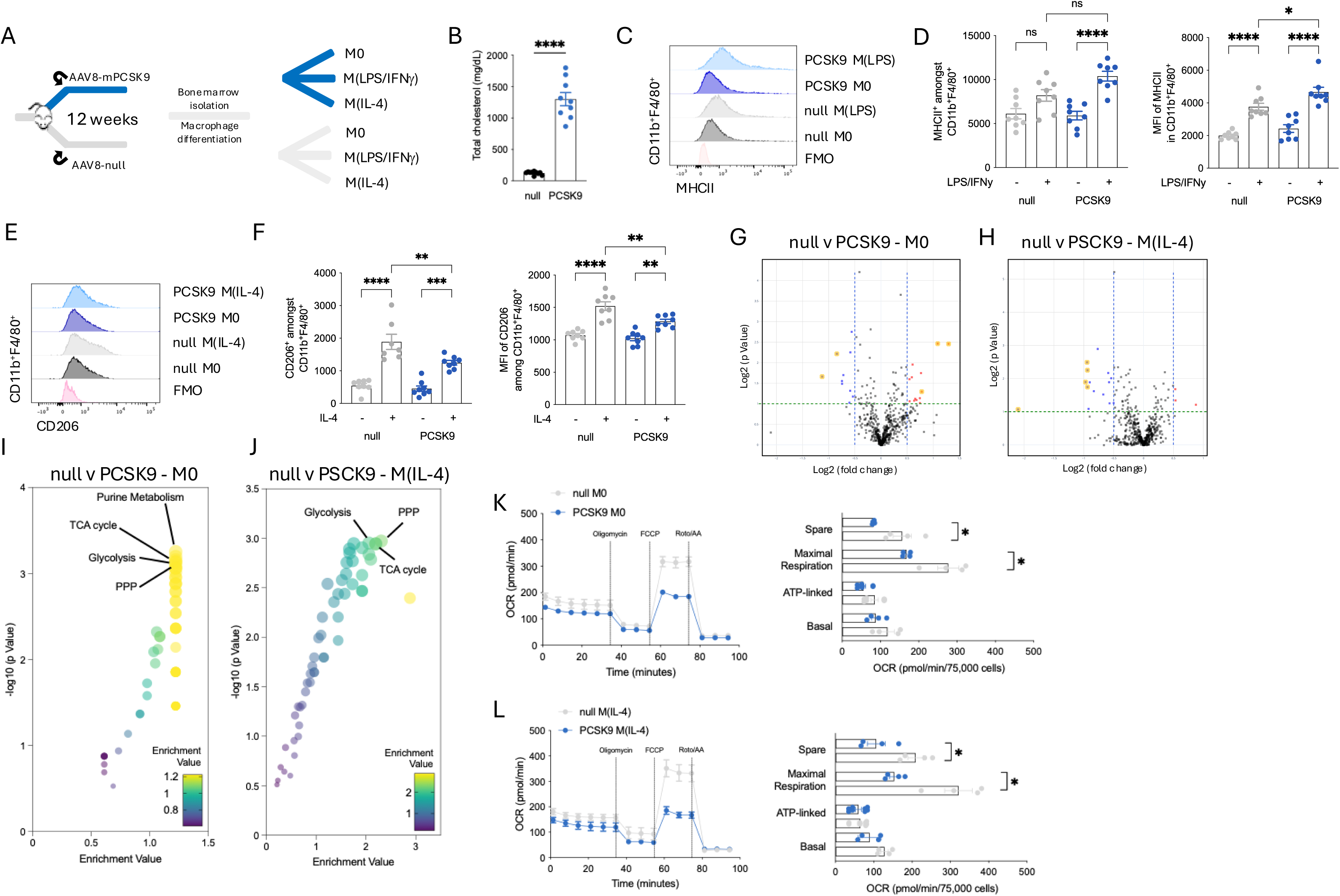
BMDM from hypercholesterolaemic mice have an altered polarisation and metabolic phenotypic with reduced resolution capacity. A. Schematic of mouse model of high cholesterol induction in C57BL/6 mice; to generate bone marrow derived macrophages. B. Plasma cholesterol level in AAV8-null and AAV8-PCSK9 mice. C. Ridge plot of MHCII expression in BMDM from AAV8-null and AAV8-PCSK9 mice.. D. Quantification of flow cytometry data for MHCII expression in M0 and M(LPS/IFNy) BMDM from AAV8-null and AAV8-PCSK9 mice. E. Ridge plot of CD206 expression in M0 and M(IL-4) BMDM from AAV8-null and AAV8-PCSK9 mice. BMDM. F. Quantification of flow cytometry data for CD206 expression in M0 and M(IL-4) BMDM from AAV8-null and AAV8-PCSK9 mice. G. Volcano plot of differential abundance of metabolites between M0 BMDM generated from AAV8-null or AAV8-PCSK9 treated mice. H. Volcano plot of differential abundance of metabolites between M(IL-4) BMDM generated from AAV8-null or AAV8-PCSK9 treated mice. I. Bubble plots of Pathway enrichment analysis of metabolomic data from M0 BMDM J. Bubble plots of Pathway enrichment analysis of metabolomic data from M(IL-4) BMDM K. Oxygen consumption rate (OCR) was measured using XFe96 Seahorse bioanalyzer with compounds used to determine basal, ATP-linked and maximum respiration in M0 BMDM generated from AAV8-null or AAV8-PCSK9 treated mice. (null M0; grey lines and PCSK9 M0; blue lines. (n = 4). L. Oxygen consumption rate (OCR) was measured using XFe96 Seahorse bioanalyzer with compounds used to determine basal, ATP-linked and maximum respiration in M(IL-4) BMDM generated from AAV8-null or AAV8-PCSK9 treated mice. (null M(IL-4); grey lines and PCSK9 M(IL-4); blue lines. (n = 4) All data are mean□±□s.e.m. of biological replicates. Significance values calculated using two-tailed Student’s *t*-test between two experimental groups one-way analysis of variance (ANOVA) for multiple comparisons, are indicated as *P<0.05, **P<0.01, ***P<0.001, ****P<0.0001.

As metabolic rewiring is a key feature of innate trained immunity ^24^, we undertook an unbiased metabolomic screen to identify key metabolic pathways which could explain the enhanced pro-inflammatory polarisation phenotype seen in BMDM from HC mice. This revealed differential abundance of metabolites related to purine metabolism and enrichment of central carbon metabolism pathways (TCA cycle, glycolysis and pentose phosphate pathway) in M0 BMDM from HC mice (Figure G/I). Similarly, M(IL-4) BMDM also had differential abundance of metabolites related to central carbon metabolism pathways (Figure 1H/J). As M(IL-4) macrophages preferentially utilise OxPhos for energy production, we first looked at the availability of pyruvate to enter into the TCA cycle. Indeed, there was a decrease in the abundance pyruvate and its precursors in BMDM from HC mice (Sup Figure 1A). Flux through the TCA cycle is essential to provide substrate for OxPhos. Early breaks in the TCA cycle were detected in both M0 BMDM (Sup Figure 1B) and M(IL-4) BMDM (Sup Figure 1C) from HC mice as evidenced by the accumulation of cis-aconitate and isocitrate (Sup Figure 1D). We hypothesised that this would reduce respiratory capacity through the mitochondrial electron transport chain. Indeed, HC-trained mice had lower oxygen consumption rates (OCR) associated with maximal respiration and spare capacity in both M0 BMDM (Figure 1K) and M(IL-4) BMDM (Figure 1L). Taken together, these results demonstrate that exposure to high cholesterol has persisting effects on myeloid cell metabolism and polarisation potential.

### The effects of HC in innate immune cells persist despite cholesterol lowering

To determine whether lipid-lowering *in vivo* could reverse the effects of exposure to high cholesterol, we administered a therapeutic anti-PCSK9 monoclonal antibody (Evolocumab) to HC mice (Figure 2A). Mice received either an injection of AAV8-mPCSK9 with HFD for 12 weeks, or null AAV8 with chow diet for the same time period. One group of HC-mice was reverted to chow diet and given a weekly subcutaneous injection of Evolocumab (denoted ‘reversal’), while the other HC group was maintained on the HFD (denoted ‘progression’; Figure 2A). There was no difference in body weight between the three groups (Figure 2B). Blood sampling confirmed that AAV8-mPCSK9 treated mice developed HC (Figure 2C). After 10 weeks, ‘progression’ mice maintained the same level of HC, but Evolocumab-treated ‘reversal’ mice had significantly lower cholesterol levels (Figure 2D).

**Figure 2:**
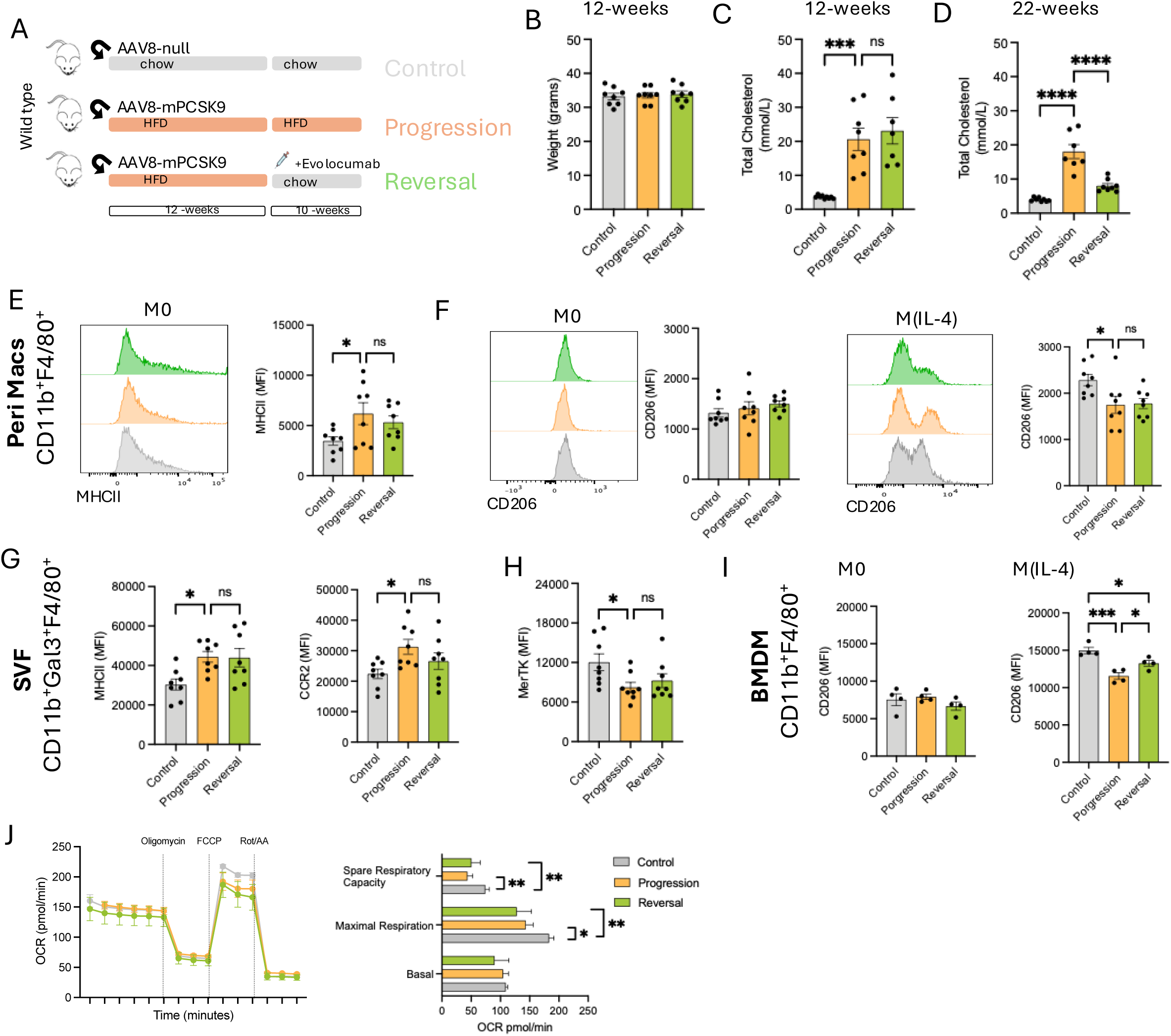
Macrophages isolated from mice following cholesterol lowering intervention retain a hyperpolarised phenotype towards an M1-like, and fail to re-polarise towards M2-like. A. Schematic of mouse model of high cholesterol induction in C57BL/6 mice with cholesterol lowering intervention using Evolocumab. B. Body weight of mice after 12 weeks on protocol. C. Total plasma cholesterol levels after 12 weeks of protocol. D. Total plasma cholesterol levels after 22 weeks of protocol. E. Flow cytometry data, mean fluorescent intensity of MHCII in CD11b+F4/80+ peritoneal macrophages stained immediately after lavage. F. Flow cytometry data, mean fluorescent intensity of CD206 in CD11b+F4/80+ peritoneal macrophages stained after 18 hour *ex vivo* M0 or after IL-4 stimulation M(Il-4). G. The stromal vascular fraction was isolated from eWAT by collagenase digestion followed by density centrifugation Flow cytometry data of mean fluorescent intensity of MHCII, CCR2 and (H) MerTK in CD11b+Gal3+F4/80+ macrophages. I. Bone marrow derived macrophages were generated and flow cytometry perform. Flow cytometry data, mean fluorescent intensity of CD206 in CD11b+F4/80+ BMDM stained after 18 hour *ex vivo* M0 or after IL-4 stimulation M(Il-4). J. Oxygen consumption rate (OCR) was measured using XFe96 Seahorse bioanalyzer with compounds used to determine basal, ATP-linked and maximum respiration in M(IL-4) BMDM generated from chow, progression and reversal mice. (Chow M(IL-4); grey lines, progression M(Il-4); orange lines and Reversal M(IL-4); green lines (n = 4)). All data are mean□±□s.e.m. of biological replicates. Significance values calculated using one-way analysis of variance (ANOVA) for multiple comparisons, are indicated as *P<0.05, **P<0.01, ***P<0.001, ****P<0.0001.

Peritoneal macrophages and adipose tissue were harvested from control, progression and reversal mice. Peritoneal macrophages from the progression group displayed a heightened pro-inflammatory phenotype, but this was not significantly altered by *in vivo* LDL-C-lowering in the reversal group (Figure 2E). Peritoneal macrophages displayed a similar inability to fully polarise when stimulated with IL-4 in both the progression and reversal groups (Figure 2F); indicating a sustained effect of exposure to HC. To determine whether these changes were also observed in tissues, adipose tissue macrophages were isolated from the stromo-vascular fraction (SVF) of HC-mice. Adipose tissue macrophages from HC mice had a pro-inflammatory phenotype (evidenced by increased MHCII and CCR2 expression), and a lower MerTK expression, indicating a reduced resolution phenotype, which persisted even after cholesterol lowering in reversal mice (Figure 2G). (Figure 2H). These data demonstrate that high cholesterol causes long-term phenotypic changes in macrophage polarisation that are sustained after cholesterol lowering and are observed in macrophages from different tissue locations.

To test whether the effect of exposure to HC in macrophages is mediated through changes in bone marrow, BMDM from each group of mice were differentiated *ex vivo*. At M0, macrophages displayed a normal polarisation profile, however, following IL-4 stimulation, progression and reversal macrophages failed to polarise into a pro-resolution type macrophage (Figure 2I). To investigate whether this effect is through long-term metabolic rewiring, we performed SeaHorse studies (Figure 2J), revealing that BMDM from HC mice have reduced spare capacity and reduced maximal OCR, which persist in reversal BMDM (Figure 2K). High content microscopy revealed no structural alternations in mitochondrial number or morphology (Sup Figure 2A; Videos Sup Figure 2B), suggesting that the sustained metabolic phenotype retained by HC-trained BMDM is not mediated by structural alternations in the mitochondrial network.

### Hematopoietic stems cells retain an epigenetic signature of exposure to HC

Since the inflammatory and metabolic effects of prior exposure to high cholesterol are maintained even in BMDM, which are derived from HSC, we next sought to determine how this effect is mediated in HSCs, by comparing bone marrow HSCs from the control, progression and reversal groups of mice. Bone marrow from mice exposed to HC had reduced long term-HSC (CD34^-^CD48^-^CD150^+^LSK) and short term-HSC (CD34^+^CD48^-^ CD150^-^LSK) populations; similar changes persisted after cholesterol lowering in the reversal mice (Figure 3A, 3B). A similar reduction in multipotent progenitor cells (MPP1) was seen in bone marrow from both progression and reversal mice (Figure 3C). In contrast, myeloid-biased MPP2 (CD48^+^CD150^-^LSK) and MMP3 (CD48^+^CD150^-^LSK) HSCs were increased in HC mice (Figure 3D), associated with a consistent increase in the overall number of myeloid cells (Figure 3E). While the proportion of MPP2/3 cells was reduced after cholesterol-lowering (Figure 3D), the total number of myeloid cells in the bone marrow did not change significantly (Figure 3E), indicating a more rapid myeloid progenitor maturation.

**Figure 3.**
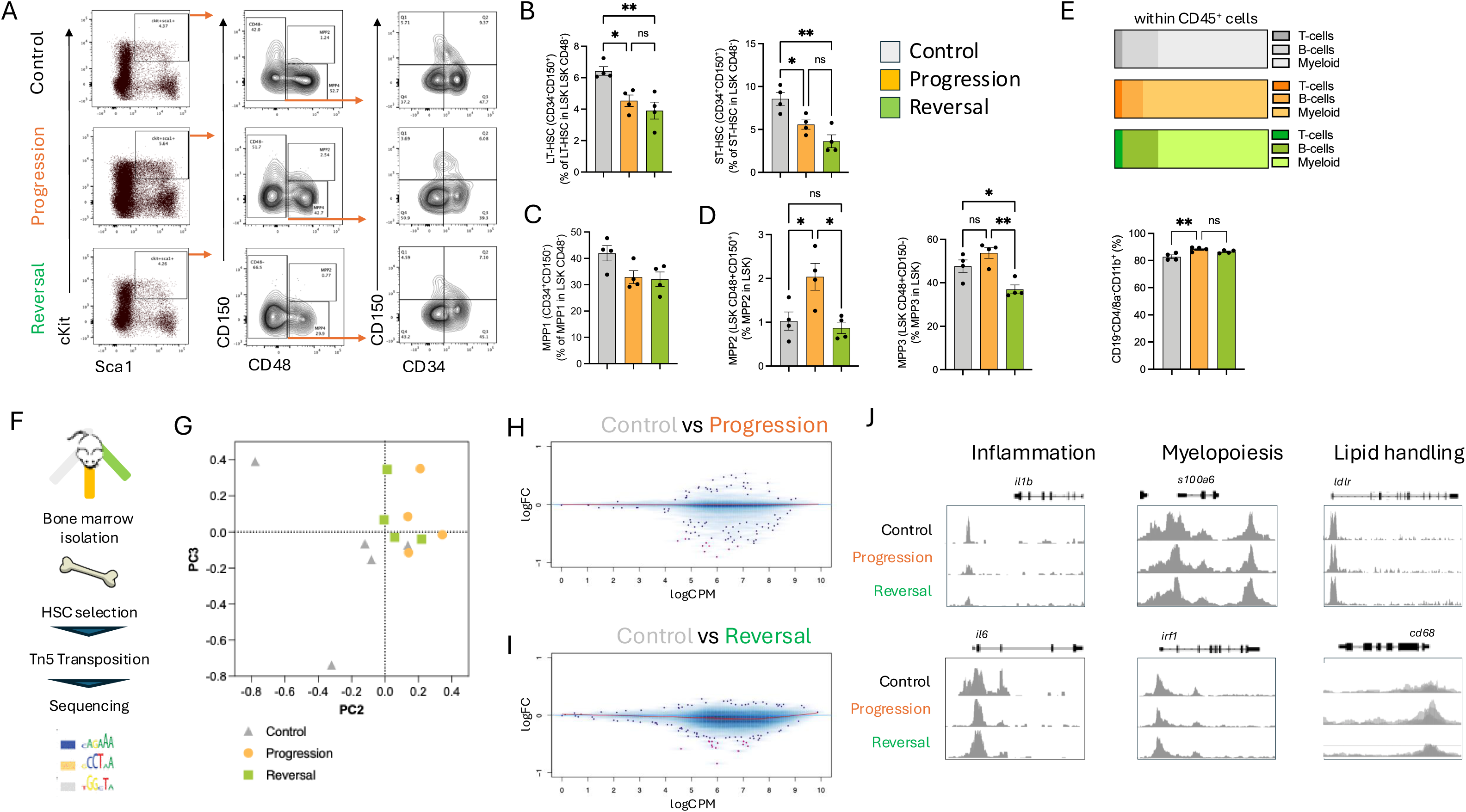
Hematopoietic stem cells (HSC) from hyperlipidemic mice have an altered phenotype which is retained even following lipid lowering A. Gating strategy for flow cytometry analysis of hematopoietic stem cells (HSC) and progenitors in the bone marrow. B. Quantification of long term and short term hematopoietic stem cells (HSC) flow cytometry in the bone marrow. C. Quantification of multi-potent progenitors 1 (MPP1) cells by flow cytometry in the bone marrow. D. Quantification of MPP2 and MPP3 cells by flow cytometry in the bone marrow. E. Quantification of mature haematological lineage cell types in the bone marrow by flow cytometry. F. Schematic of experimental work flow of assessing chromatin accessibility using ATAC-Seq ATAC-Seq of 500 lin-ckit+Sca1+ hematopoietic stem cells. G. Principal Component Analysis (PCA) plot called peak following ATAC-Seq of lin-ckit+Sca1+ hematopoietic stem cells. H. MA plot for transposase accessible chromatin regions between chow and progression lin-ckit+Sca1+ hematopoietic stem cells I. MA plot for transposase accessible chromatin regions between chow and reversal lin-ckit+Sca1+ hematopoietic stem cells J. Merged Integrative Genome Browser track plots for chow, progression and reversal ATAC-Seq of lin-ckit+Sca1+ hematopoietic stem cells, for genes related to inflammation (*il1b* and *il6*), myelopoiesis (*s100a6* and *irf1*) and lipid handling (*ldlr* and *cd68*). All data are mean□±□s.e.m. of biological replicates. Significance values calculated using one-way analysis of variance (ANOVA) for multiple comparisons, are indicated as *P<0.05, **P<0.01, ***P<0.001, ****P<0.0001.

ATAC-sequencing (ATAC-seq) of Lin^-^Sca1^+^cKIT^+^ HSC (Figure 3F) revealed that exposure to HC results in wide-ranging changes to the chromatin landscape that persisted were not fully reversed after cholesterol-lowering (Figure 3G). Significant differences in ATAC-seq peaks were detected between control vs. progression (Figure 3H) and control vs. reversal HSCs (Figure 3I) but no difference in ATAC-seq peaks were detected between progression and reversal HSCs at 5 % FDR. The differential chromatin accessibility between control and HC HSC included regions related to proinflammatory cytokine production (*il1b*, *il6*), myelopoiesis (*irf1*, *s100a6*) and lipid handling (l*dlr*, *cd68*) (Figure 3J). Importantly, chromatin accessibility in HSCs from reversal mice remained significantly different to that in control HSC, indicating that the changes in chromatin landscape are maintained after cholesterol lowering (Figure 3I/J.

### High cholesterol induces chromatin remodelling and altered Runx1-dependent regulation of fatty acid metabolism to reduce macrophage M2 polarisation

To identify the molecular and metabolic basis for the effect of prior HC exposure on macrophage polarisation, we evaluated chromatin accessibility in bone marrow monocytes, the precursors of tissue resident macrophages (Figure 4A). Differential chromatin accessibility was identified in a total of 2794 regions with an FDR < 1 %. Of the 1376 regions with increased accessibility, 21 had a >2-fold change. These included regions in close proximity to genes including *Tubb1*, *Cd226*, and *Scd1* (Figure 4B). KEGG pathway analysis of the genes nearest to differentially accessible peaks highlighted pathways involved in immune system process, cholesterol handling inflammatory response and mono-unsaturated fatty acid (MUFA) biosynthesis (Figure 4C). Genes related to mono-unsaturated fatty acid (MUFA) biosynthesis (*Scd1*, *Scd2*, *Scd3* and *Scd4*), all exhibited increased chromatin accessibility within promoter regions (Figure 4D; Sup Figure 3A). This was associated with decreased expression of *Scd1* in HC-trained BM monocytes (Figure 4E). As SCD is the rate limiting step in the production of MUFA, we hypothesised that HC-trained BMDM would have lower concentrations of MUFA, Indeed, metabolomics analysis revealed that MUFA abundance was decreased (Figure 4F) between HC-trained and control M(IL-4) BMDM, while other lipid species including sterols remained unaltered (Sup Figure 3B).

**Figure 4:**
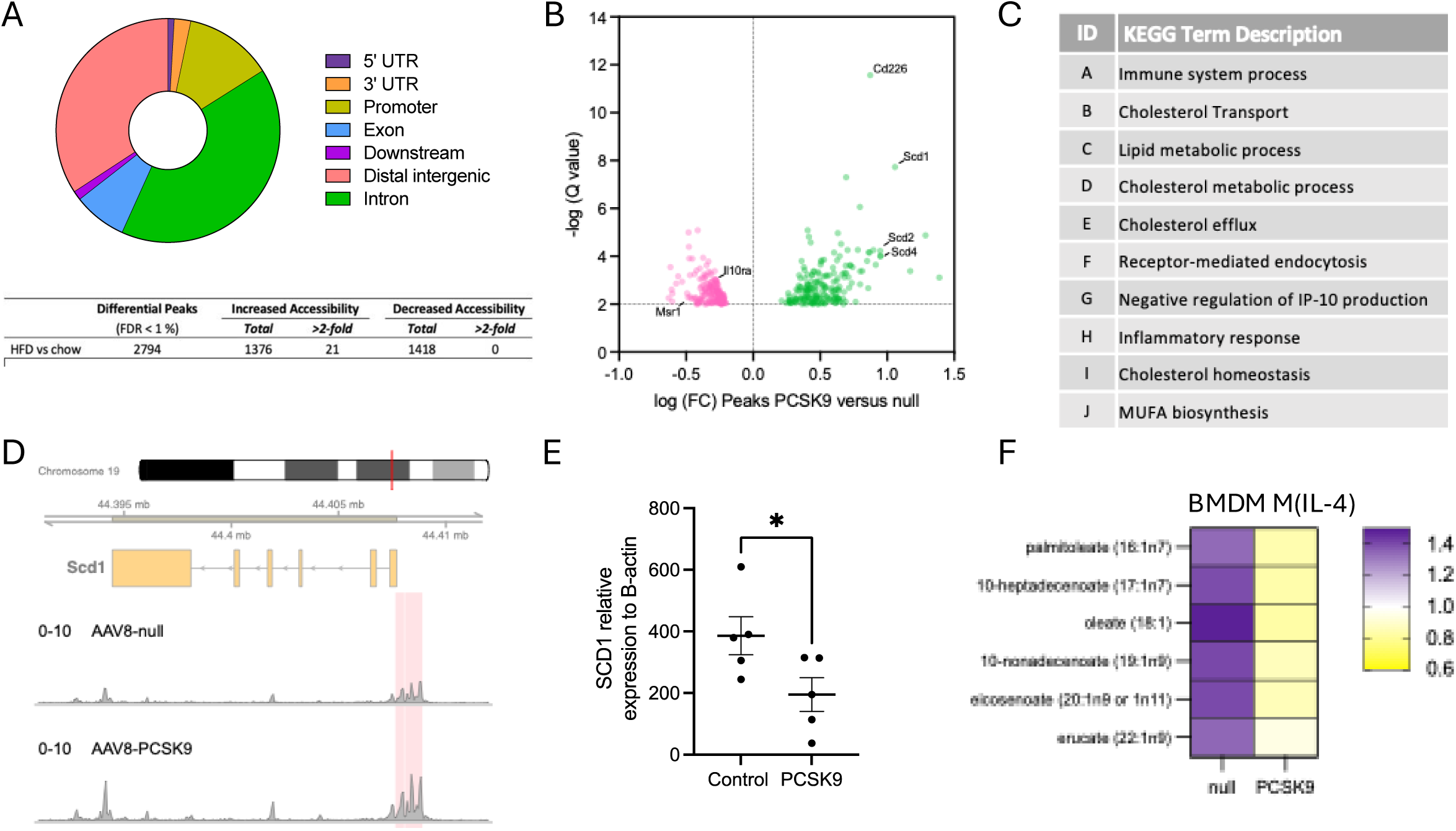
HC-induced changes to SCD alter monocytes and macrophage MUFA abundance. A. Doughnut plot of location of peaks, and table denoting directionality of peaks which read 1 % FDR of ATAC-Seq data in bone marrow monocytes from AAV-null and AAV8-PCSK9 mice. B. Volcano plot of differentially accessible peaks between bone marrow monocytes isolated from AAV-null and AAV8-PCSK9 mice. C. Top 10 enriched KEGG pathways base on the top 250 peaks identified in ATAC-seq in bone marrow monocytes from AAV-null and AAV8-PCSK9 mice D. Scd1 gene loci showing merged Integrative Genome Browser tracks of ATAC-seq peaks. ATAC-seq peaks shaded in red are differentially accessible in bone marrow monocytes from AAV-null and AAV8-PCSK9 mice E. Relative gene expression of Scd1 in bone marrow monocytes from AAV-null and AAV8-PCSK9 mice. F. Relative abundance of mono-unsaturated fatty acids in M(IL-4) BMDM generated from AAV-null and AAV8-PCSK9 mice. Data in (E) are mean□±□s.e.m. of biological replicates. Significance values calculated using Student’s t-test, are indicated as *P<0.05.

### MUFA metabolism mediates changes in macrophage polarisation

To test whether MUFA production via SCD is important for macrophage polarisation in response to HC, fully differentiated BMDM were exposed to an SCD inhibitor during polarisation (Sup Figure 3C). SCD inhibition had no effect on polarisation (Sup Figure 3D), or cellular metabolism (Sup Figure 3E) in differentiated BMDM. In contrast, treatment of BM with an SCD inhibitor (Figure 5A) throughout BMDM differentiation resulted in an exaggerated pro-inflammatory phenotype in the resultant M(IL-4) BMDM (Figure 5B) and reduced OxPhos capacity (Figure 5C/D), recapitulating the effect of prior HC-exposure. We next sought to investigate whether oleic acid (OA) supplementation to BMDM could enhance OxPhos capacity by providing extra substrate for fatty acid oxidation and thus promoting the M(IL-4) phenotype. Cells treated with OA during M(IL-4) polarisation had increased cell surface expression of CD206 (Figure 6E), and increased basal, maximum and spare OCR (Figure 5F-I), suggesting that supplementation with exogeneous OA enhances OxPhos. Consistent with this, the increase in spare capacity OCR seen in OA-treated M(IL-4) BMDM was reduced by treatment with Etomoxir, which inhibits mitochondrial FA uptake (Figure 5J-M).

**Figure 5:**
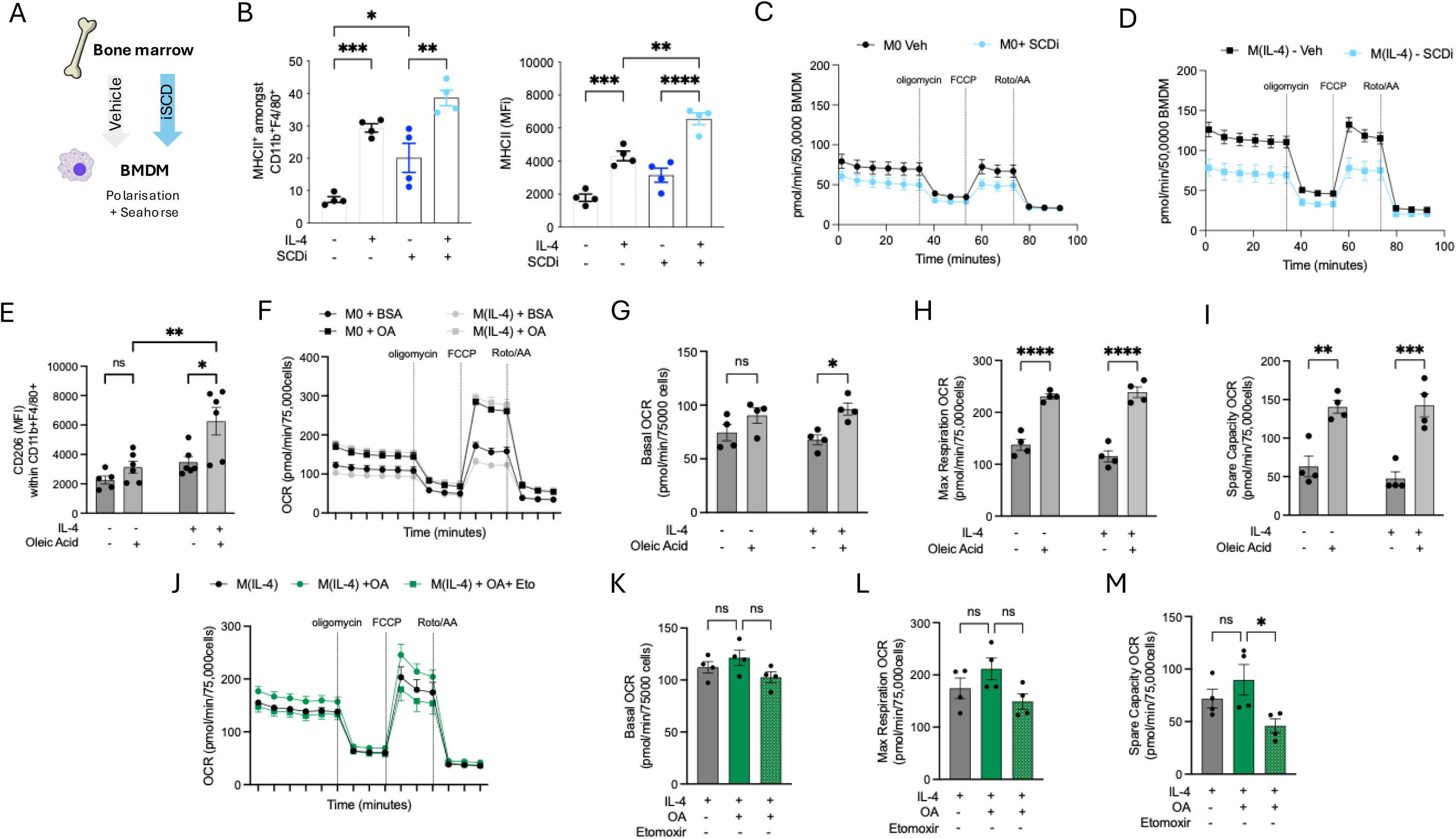
Role of MUFA in M(IL-4) macrophage metabolism and polarisation. A. Schematic of bone marrow derived macrophage (BMDM) experiment with SCD inhibition. B. Histogram showing quantification of MHCII expression amongst CD11b+F4/80+ BMDM treated with SCDi during BM to BMDM. C. Oxygen consumption rate (OCR) was measured using XFe96 Seahorse bioanalyzer with compounds used to determine basal, ATP-linked and maximum respiration in M0 BMDM treated with SCDi during BM to BMDM differentiation. (M0 Vehicle; black lines and M0 + SCDi; pale blue lines. (n = 4). D. Oxygen consumption rate (OCR) was measured using XFe96 Seahorse bioanalyzer with compounds used to determine basal, ATP-linked and maximum respiration in M(Il-4) BMDM treated with SCDi during BM to BMDM differentiation. (M(Il-4) + Vehicle; black lines and M(IL-4) + SCDi; pale blue line. (n = 4). E. Relative expression of CD206 on BMDM treated with Oleic acid during M(IL-4) polarisation. F. Oxygen consumption rate (OCR) was measured using XFe96 Seahorse bioanalyzer with compounds used to determine basal, ATP-linked and maximum respiration in M0 and M(IL-4) BMDM treated with Oleic acid during polarisation. (M0 + BSA; circles black line; M0 + oleic acid; squares black line; M(IL-4) + BSA; circles grey line and M(IL-4) + oleic acid; squares grey line. (n = 4). G. Quantified basal oxygen consumption rate from (G) H. Quantified maximal respiration oxygen consumption rate from (G).. I. Quantified spare capacity oxygen consumption rate from (G). J. Oxygen consumption rate (OCR) was measured using XFe96 Seahorse bioanalyzer with compounds used to determine basal, ATP-linked and maximum respiration in M(IL-4) BMDM treated with Oleic acid in the presence of Etomoxir during polarisation. (M(IL-4) black line; M(IL-4) + oleic acid circles green line; M(IL-4) + oleic acid + etomoxir squared green line (n = 4). K. Quantified basal oxygen consumption rate from (K). L. maximal respiration oxygen consumption rate from (K). M. Quantified spare capacity oxygen consumption rate from (K). All data are mean□±□s.e.m. of biological replicates. Significance values were calculated using one-way analysis of variance (ANOVA) for multiple comparisons, are indicated as *P<0.05, **P<0.01, ***P<0.001, ****P<0.0001.

**Figure 6:**
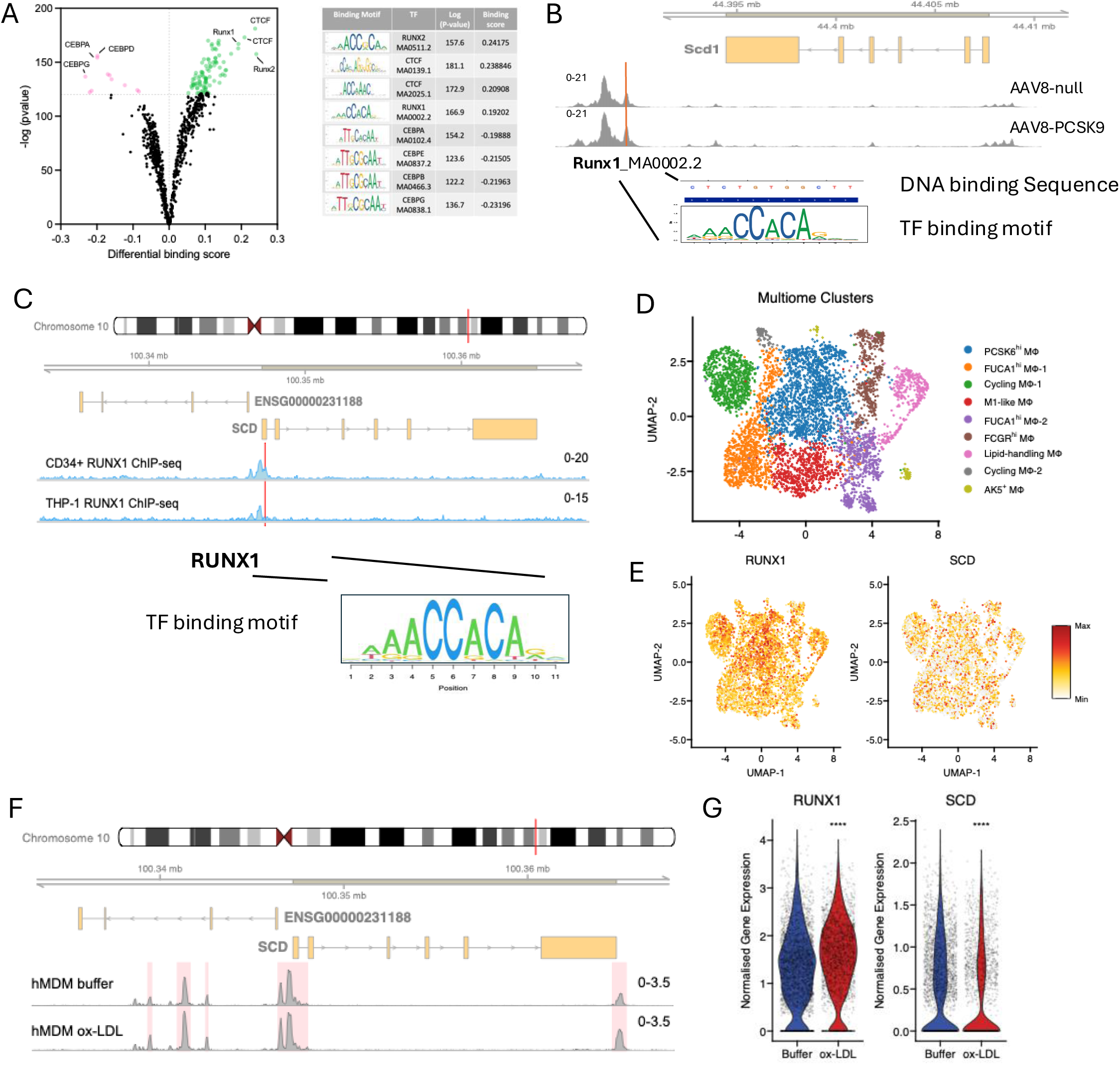
Runx1 mediates the cholesterol-induced dysregulation of SCD A. Volcano plot of transcription factor binding motifs.. Top 8 transcription factor motifs identified in differentially enriched ATAC-seq peaks in bone marrow monocytes from AAV-null and AAV8-PCSK9 mice. B. Scd1 gene loci showing merged Integrative Genome Browser tracks of ATAC-seq peaks. Identified is a predicted RunX1 TF bind motif with in differentially accessible peaks from ATAC-seq data in bone marrow monocytes from AAV-null and AAV8-PCSK9 mice. C. Human Scd gene loci showing merged Integrative Genome Browser tracks of CD34+ RUNX1 CHIP-Seq data and THP-1 RUNX1 CHIP-seq data. Red line shows the location of predicted RUNX1 transcription factor binding motif. D. UMAP projection of single nuclei multiome data of human monocyte derived macrophages treated with oxLDL or buffer. Individual clusters coloured. E. Relative expression of RUNX1 and SCD shown in UMAP projection. Expressed as minimum and max projections. F. Scd gene loci showing merged Integrative Genome Browser tracks of human monocyte derived macrophages treated with oxLDL vs buffer. Red boxed indicate significant peaks. G. Normalised gene expression of RUNX1 and SCD in human monocyte derived macrophages treated with oxLDL or buffer.

### Runx1 mediates the cholesterol-induced dysregulation of SCD

Next, we aimed to characterise the transcriptional regulators responsible for the changes in chromatin accessibility in HC-trained monocytes. Motif footprinting analysis revealed an increased binding for transcription factors such as CTCF, Runx1, and Runx2, and a decreased binding for members of the CEBP family (Figure 6A). We identified a putative Runx1 binding site 5kb downstream of the Scd1 gene, with a concordant increase in accessibility in the Scd1 promoter (Figure 6B). In addition, RUNX1 ChIP-seq in human CD34+ HSC and monocytic THP-1 cells revealed significant binding of RUNX1 at the promoter of the SCD gene, which is homologous to the mouse Scd genes (Figure 6C). This finding confirms that the regulatory relationship between RUNX1 and SCD is conserved in both mice and humans. Next, we explored whether exposure to cholesterol in human monocyte-derived macrophages (hMoDM) would yield a similar effect on SCD, mediated by RUNX1. In Single-cell multiome data in hMoDM we identified 9 transcriptionally distinct clusters (Figure 6D/E), we found that RUNX1 and SCD were widely expressed in all ^14^. The SCD promoter region became significantly more accessible after exposure to ox-LDL (Figure 6F). Simultaneous gene expression measurements in the same cells showed an increase in RUNX1 expression, along with a decrease in SCD expression (Figure 6G). This pattern of decreased SCD expression despite increased promoter accessibility was consistent in both HC-trained mouse monocytes and ox-LDL-exposed hMoDM. Taken together, these observations suggest that epigenetic changes in HSC, and myeloid progenitors, lead to altered expression of key enzymes involved in MUFA production, promoting a trained pro-inflammatory phenotype, by restricting the metabolic plasticity of macrophages to polarise in to a resolution phenotype.

### Prior exposure to high cholesterol promotes pro-inflammatory macrophage polarisation following bone marrow transplantation

As exposure to HC promotes long-term metabolic reprograming in BM HSCs with skewing of myeloid cell differentiation towards a pro-inflammatory state, we reasoned that this phenotype should persist in recipient mice following bone marrow transplantation. To test this hypothesis, we isolated BM from HC or control mice expressing GFP under the control of the human CD68 promoter (hCD68-GFP mice) in order to identify fluorescent donor-derived myeloid cells. BM from either control or HC hCD68-GFP mice was transplanted into irradiated normocholesterolaemic recipient wild-type mice (Figure 7A). As expected, hCD68 mice treated with AAV8-mPCSK9 and fed a HFD had significantly elevated plasma cholesterol levels and no difference in weight (Figure 7B). Recipient mice after BM transplantation had physiological LDL-cholesterol levels, irrespective of whether the transplanted BM was derived from HC hCD68-GFP donors (Figure 7C). Chimerism was confirmed by quantifying expression of hCD68-GFP in circulating neutrophils (∼90 %; Sup Figure 5A/B), while non-myeloid cells expressed less <5 % (Sup Figure 4A/C). Peritoneal macrophages were isolated from chimeric mice and cultured *ex vivo*. By 7 weeks after BMT, mice which received HC donor BM had increased repopulation of the peritoneum with hCD68 GFP+ cells. (Figure 7D). M0 peritoneal macrophages displayed no differences in expression of MHCII (Figure 7E), but CD206 expression was reduced in peritoneal macrophages in mice transplanted with HC donor BM (Figure 7F). When peritoneal macrophages were stimulated with LPS/IFNg there was further enhancement of the pro-inflammatory phenotype in mice that had received BM from a HC-donor (Figure 7G). Furthermore, peritoneal macrophages failed to polarise towards a pro-resolution-type macrophage when stimulated with IL-4, if the host mice had received BM from a HC-donor (Figure 7H). BMDM from HC donor mice had increased TNFa and IL-6 secretion when stimulated with LPS/IFNg (Sup Figure 4D), indicating that they are primed to have an exaggerated pro-inflammatory response. Critically, BMDM generated from chimeric mice retained the same metabolic capacity as the HC-donor mice, i.e. reduced spare capacity and maximal respiration OCR in both M0 BMDM (Figure 7I) and M(IL-4) BMDM (Figure 7J). These observations demonstrate that the effects of HC *in vivo* are retained after BM transplantation and mediated by epigenetic changes in HSC progenitors and their derived myeloid cells, that alter metabolism and are refractory to the reduction of cholesterol in the recipient mouse.

**Figure 7:**
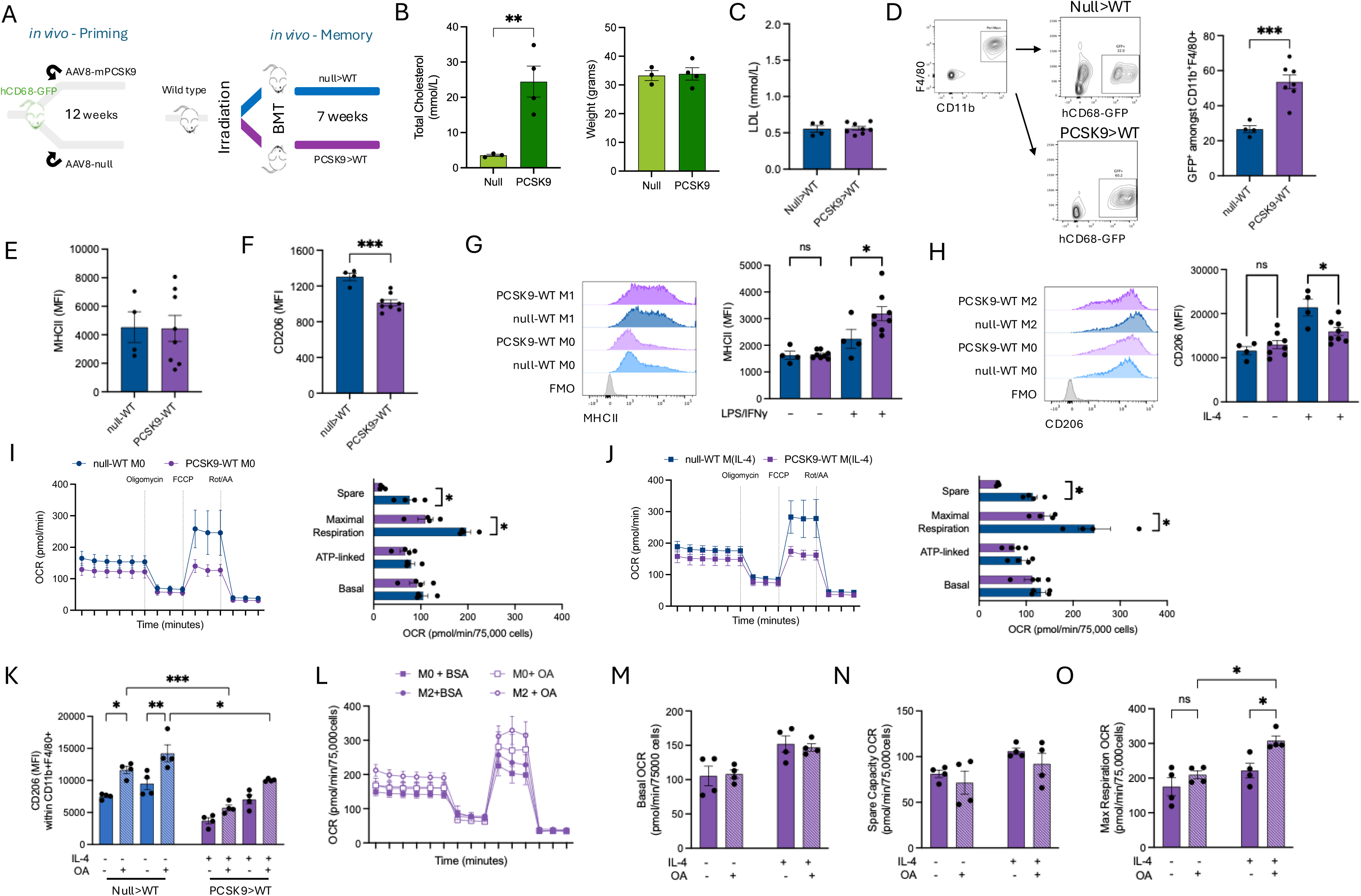
BMDM grown from bone marrow transplanted hyperlipidemic mice retained an altered phenotype even after systemic cholesterol is normalised. A. Schematic of mouse model of bone marrow transplant of high cholesterol-trained bone marrow or normal cholesterol bone marrow from hCD68-GFP mice into irradiated C57BL/6 mice. B. Body weight and total plasma cholesterol level in bone marrow transplant mice. C. Plasms LDL cholesterol levels in recipient mice after 7 weeks. D. Flow cytometry data, & of hCD68-GFP+ cells within CD11b+F4/80+ peritoneal macrophages. E. Flow cytometry data, mean fluorescent intensity of MHCII in CD11b+F4/80+ peritoneal macrophages stained immediately after lavage. F. Flow cytometry data, mean fluorescent intensity of CD206 in CD11b+F4/80+ peritoneal macrophages stained immediately after lavage. G. Flow cytometry data, mean fluorescent intensity of MHCII in CD11b+F4/80+ peritoneal macrophages stimulated with LPS/IFNy for 18 hours. H. Flow cytometry data, mean fluorescent intensity of CD206 in CD11b+F4/80+ peritoneal macrophages stimulated with LPS/IFNy for 18 hours. I. Oxygen consumption rate (OCR) was measured using XFe96 Seahorse bioanalyzer with compounds used to determine basal, ATP-linked and maximum respiration in M0 BMDM (null>WT M0; blue lines and PCSK9>WT M; purple line. (n = 4). With quantification of spare capacity, maximal respiration, ATP-link and basal OCR. J. Oxygen consumption rate (OCR) was measured using XFe96 Seahorse bioanalyzer with compounds used to determine basal, ATP-linked and maximum respiration in M(Il-4) BMDM (null>WT M(IL-4); blue lines and PCSK9>WT M(IL-4); purple line. (n = 4). With quantification of spare capacity, maximal respiration, ATP-link and basal OCR. K. Flow cytometry data, mean fluorescent intensity of CD206 in CD11b+F4/80+ BMDM supplemented with LPS/IFNy for 18 hours. L. Oxygen consumption rate (OCR) was measured using XFe96 Seahorse bioanalyzer with compounds used to determine basal, ATP-linked and maximum respiration in M(Il-4) BMDM treated with Oleic acid or BSA. null>WT M(IL-4 + BSA), Solid purple square; (null>WT M(IL-4 + OA), Open purple square; (PCSK9>WT M(IL-4 + BSA), Solid purple circle and (PCSK9>WT M(IL-4 + OA), Open purple circle. (n = 4). M. Quantified basal oxygen consumption rate from (L). N. Quantified spare capacity oxygen consumption rate from (L). O. Quantified maximal respiration oxygen consumption rate from (L). All data are mean□±□s.e.m. of biological replicates. Significance values calculated using two-tailed Student’s *t*-test between two experimental groups one-way analysis of variance (ANOVA) for multiple comparisons, are indicated as *P<0.05, **P<0.01, ***P<0.001, ****P<0.0001.

We next tested whether the reduced metabolic plasticity in BMDM generated from mice transplanted with HC-donor BM could be overcome by MUFA supplementation, in order to provide additional substrate for OxPhos, that might promote M(IL-4) polarisation. Oleic acid (OA) treatment of BMDM from control chimeric mice increased cell surface expression of CD206, indicating an enhanced M2-like phenotype (Figure 7K). BMDM generated from mice transplanted with HC-donor BM also showed increases in CD206 expression after OA treatment (Figure 7K). When OCR was assessed in M(IL-4) BMDM OA supplementation did not increase basal or spare capacity OCR, but did increase maximal respiration capacity (Figure 7L-O) in BMDM generated from mice transplanted with HC-donor BM. These results demonstrate that OA supplementation can increase OxPhos capacity and restore the polarisation potential of HC-trained macrophage.

### HC-exposed bone marrow confers systemic metabolic dysfunction through enhanced repopulation of adipose tissue macrophages with increased pro-inflammatory polarisation

Since myeloid cell inflammation is associated with adverse cardiometabolic outcomes, we reasoned that HC-trained bone marrow would lead to repopulation of tissues and organs with macrophages of a pro-inflammatory phenotype, with systemic metabolic consequences secondary to increased adipose tissue inflammation. Notably, chimeric mice receiving BMT from HC donor mice gained significantly more weight (Figure 8A). The SVF of adipose tissue had similar total number of adipose tissue macrophages in all transplanted mice (Figure 8B/C), which displayed similar polarisation phenotype (Supp Figure 6A). However, the SVF of mice transplanted from HC donors had significantly increased numbers and proportions of hCD68-GFP+ cells (Figure 8D), indicating that myeloid cell recruitment to adipose tissue is increased in mice receiving BM from HC donors, driven by cells derived from HSCs exposed to prior HC.

**Figure 8:**
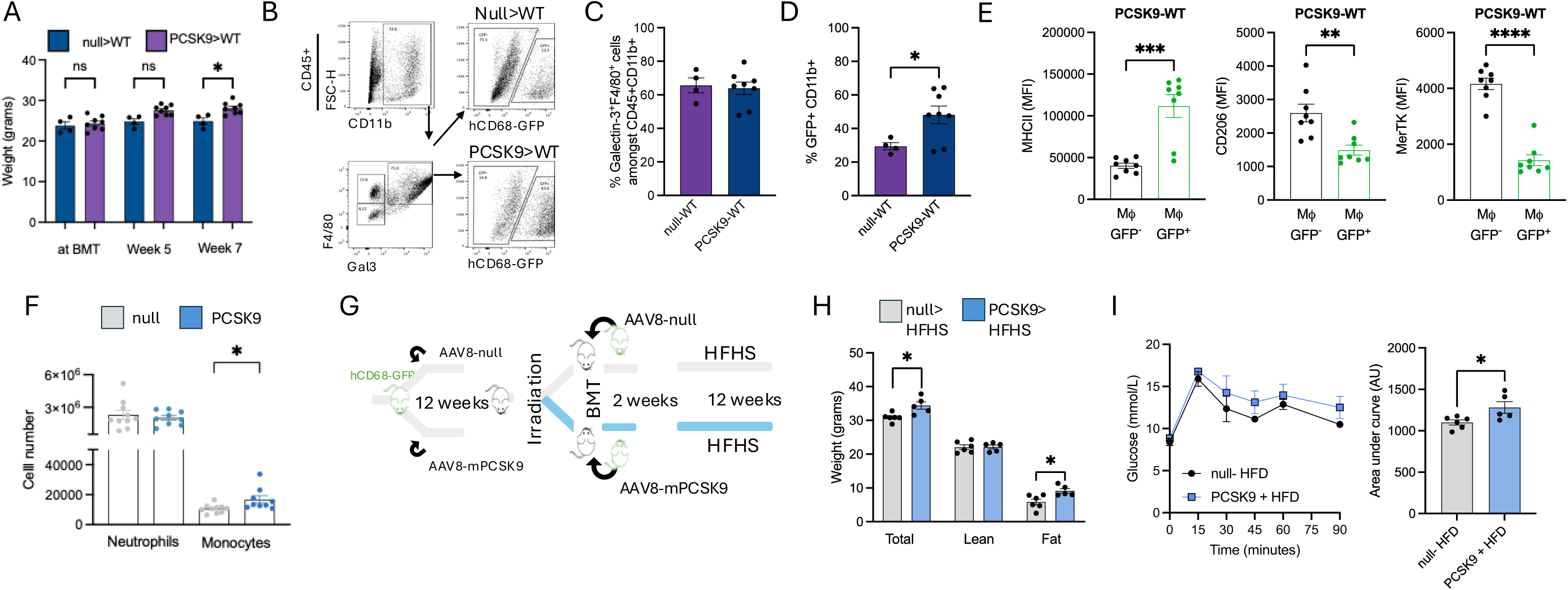
BMT of hyperlipidemic BM confers a global metabolic phenotype to recipient mice, increasing their capacity to gain weight following HFD feeding directed by increased adipose tissue inflammation. A. Weight gain following bone marrow transfer in null>WT and PCSK9>WT chimeric mice. B. Flow cytometry gating strategy of stromal vascular fraction showing hCD68-GFP expression in adipose tissue macrophages. C. Flow cytometry data quantifying the number of adipose tissue macrophages in the adipose stromal vascular fraction. D. Flow cytometry data quantifying the number of hCD68-GFP cells within the myeloid compartment of the adipose stromal vascular fraction. E. Flow cytometry data demonstrating mean fluorescent intensity of MHCII, CD206 and MerTk in GFP- and GFP+ macrophages in the adipose stromal vascular fraction of PCSK9>WT chimeric mice. F. Flow cytometry of data of zymosan induced peritonitis model. Quantification of neutrophils (CD11b+Ly6G+Ly6C+) and monocytes (CD11b+Ly6G-Ly6C+). G. Schematic of mouse model of high cholesterol induction in hCD68-GFP mice, followed by bone marrow transfer into irradiated WT mice; followed by feeding a high fat high sugar diet. H. Quantification of body composition by DEX-MRI; total body weight, lean mass and fat mass. I. Oral glucose tolerance test in fast mice and quantification of area under the curve (AUC). All data are mean□±□s.e.m. of biological replicates. Significance values calculated using two-tailed Student’s *t*-test between two experimental groups one-way analysis of variance (ANOVA) for multiple comparisons, are indicated as *P<0.05, **P<0.01, ***P<0.001, ****P<0.0001.

We next compared the polarisation state of both recipient-derived cells (GFP-) and donor-derived cells (GFP+) in the SVF. Chimeric mice receiving BM from control GFP+ donors displayed no difference in macrophage markers between GFP- and GFP+ macrophages (Sup Figure 5B). In contrast, GFP+ adipose tissue macrophages from chimeric mice receiving BM from HC GFP+ donors displayed marked increases in expression of pro-inflammatory markers and decreased homeostatic markers compared to the recipient-derived cells (GFP-) (Figure 8E), indicating that macrophages derived from HC-trained BM develop a polarisation phenotype that is maintained in adipose tissue similar to that of macrophages exposed to HC. This pro-inflammatory phenotype was further evidenced at tissue levels by increased mRNA levels of *tnfa, il6, il1b, ccl2* and *vcam* in the adipose tissue of mice receiving BM from HC donors (Sup Figure 5C). Furthermore, monocyte recruitment to the peritoneum after zymosan challenge was also increased in mice receiving BM from HC donors (Figure 8F), demonstrating that HC-trained BM are primed for recruitment.

Since adipose tissue inflammation is a driver of systemic metabolic dysfunction, we next hypothesized that the increased number and pro-inflammatory polarisation of macrophages recruited to adipose tissue could promote a metabolic syndrome phenotype. To test this hypothesis, chimeric mice received BM from either control or HC mice and were subsequently fed with a high-fat high-sugar (HFHS) diet for 12-weeks (Figure 8G). Mice which received HC-trained BM transplant gained more weight, driven by a selective increase in body fat (Figure 8H). Following an oral glucose tolerance test, mice which received HC-donor BM displayed significantly higher and prolonged increases in blood glucose levels (Figure 8I), indicating systemic glucose intolerance. These results demonstrate for the first time that HC-exposure confers a systemic metabolic effect on recipient mice beyond altering myeloid cell phenotype. These data provide evidence that long-term epigenetic consequences of HSC exposure to high cholesterol lead to an increase in risk factors for cardiometabolic disease beyond those seen in the vasculature.

## Discussion

Although residual risk in ASCVD after cholesterol-lowering is associated with inflammation, the mechanisms mediating these effects, and the rational basis for new therapeutic interventions, are not well understood. We demonstrate that HC causes aberrantly trained innate immunity through long-term effects on HSC. This results in skewed myeloid populations with reduced metabolic plasticity which favours a pro-inflammatory macrophage phenotype. Remarkably, these effects are maintained even after cholesterol-lowering, due to epigenetic reprograming that reduces macrophage OxPhos capacity. This reduction arises from changes in central carbon metabolism, reduced flux through the TCA cycle and reduced MUFA abundance, mediated by Runx1-dependent downregulation of stearoyl-CoA desaturase. HC-induced trained immunity results in increased recruitment of pro-inflammatory macrophages to adipose tissue, increased adipose inflammation and adverse systemic metabolic effects. Collectively, these results demonstrate that exposure to HC has persistent and wide-reaching effects that are maintained after cholesterol-lowering and driven through epigenetic reprogramming of HSC, highlighting the need for new approaches to reduce residual risk in ASCVD, beyond further cholesterol-lowering.

Macrophage polarisation state is intrinsically linked to metabolic plasticity^25^ that allows macrophages to respond to external stimuli; or at basal state maintain tissue homeostasis. Previous studies reported that murine macrophages exposed to high cholesterol have decreased OxPhos capacity through a defective pentose phosphate pathway^26^. We now demonstrate that this reduction in OxPhos capacity in mature macrophage is a result of epigenetic programming in HSC, and is maintained over multiple rounds of cell division as HSC divide and mature to become macrophage. Furthermore, we demonstrate that this metabolic dysfunction is maintained after LCL-C lowering. This is the case whether when cholesterol is lowered by pharmacologic and dietary intervention in HC animals, or by transplantation of BM from HC to normocholesterolemic animals. Reduced OxPhos capacity results in macrophage skewing to a pro-inflammatory phenotype, and an inability to fully polarise towards a homeostatic M2-like cell. Having intact OxPhos in monocytes/macrophages is essential for successful tissue recruitment ^14^, maintenance of their tissue homeostatic functions 15, and polarisation towards a resolution-type phenotype^27^.

Central to these new observations are changes in bone marrow HSC in response to HC exposure, that persist after cholesterol-lowering. We show that HC changes the composition of the BM HSC compartment, reducing LT-HSC number, increasing the number of multi-potent progenitors, and that after cholesterol-lowering both LT and ST-HSC number remain reduced. These observations accord with findings of reduced LT-HSC number in LDL-/-mice fed an intermittent high fat diet ^11^, suggesting a common effect of ‘harm’ to LT-HSC survival. These studies provide further evidence that stem cells niches are sensitive to changes in lipid burden, as indicated by previous studies showing that cholesterol accumulation in pluripotent HSC (PHSC) is associated with skewing of haematopoiesis toward the myeloid lineage ^28^ and that fatty acid oxidation drives asymmetric division of PHSC populations ^29^. We now demonstrate that, beyond changes to HSC number and bias, HSC undergo wide-ranging chromatin remodelling in response to HC which is not fully reversed by cholesterol-lowering. These epigenetic changes are maintained after BM transplantation, as macrophages from HC-trained BM chimeric mice with normal cholesterol levels retain the same metabolic and polarisation phenotypes as macrophages from mice with HC.

We have identified a novel mechanism linking the effects of high cholesterol to programmed changes in HSC by identifying a cluster of genes with altered chromatin accessibility following HC-training in both HSC and bone marrow monocytes. These genes are related to inflammatory function and fatty acid/lipid metabolism, including SCD which encodes a critical regulator of the production of mono-unsaturated fatty acid (MUFA), that can be oxidised to provide a substrate for OxPhos30. Our integrative analyses highlight the role of Runx1 in regulating SCD genes in both murine and human monocytes/macrophages in response to HC. Runx1 has well characterized roles in the regulation of haematopoiesis. Patients with familial hypercholesterolaemia have reduced SCD expression in PBMC compared with matched normo-cholesterolaemic controls ^31^ providing evidence that the pathway we identified could be important in understanding myeloid cells metabolic plasticity in response to cholesterol exposure.

Runx1 deficiency promotes M2 polarisation in macrophages32 and promotes myeloid bias in GMP Runx1-/-mice, both of which are key feature in HC-induced trained immunity. We identified a non-reversible myeloid bias in HSC’s, consistent with Lavillegrand et al. who reported that Runx1 is a regulator of myelopoiesis following HFD-re-exposure ^11^. Furthermore, we provide proof-of-concept evidence that pharmacological inhibition of SCD induces a phenotype similar to HC-training and that supplementation of oleic acid as an OxPhos substrate to HC-trained macrophages reverses the effects on polarisation. Blocking FA uptake into the mitochondria attenuated the beneficial effects of OA on OxPhos capacity. Thus, our work identifies Runx1-dependent regulation of fatty acid metabolism as a new, potentially targetable mechanism that mediates the programmed effects of HC on innate immune cells.

Other models of CVD-risk factor induced trained immunity have focused on progression of atherosclerotic plaques or the role of HFD/obesity-induced innate trained immunity ^9,11,12^. However, as all tissues are populated with macrophages derived from HSC, non-vascular tissues could retain a memory of previous exposure to HFD/obesity ^8,33^. Accordingly, we show that the programmed effects of HC that are refractory to cholesterol-lowering exert adverse systemic effects in other tissues. Adipose tissue inflammation is associated with obesity-related inflammation and increased immune cell infiltration ^34^. We show that HC-training increases both recruitment and pro-inflammatory polarisation of adipose tissue macrophages, which drives systemic metabolic dysfunction. This shift in adipose tissue inflammation causes an increase in body fat mass, and systemic glucose intolerance, both of which are independent risk factors for ASCVD.

In summary, we provide evidence that hypercholesterolaemia results in long-term immuno-metabolic consequences that persist despite optimal cholesterol-lowering and promote systemic inflammation and metabolic dysfunction. These effects are mediated through epigenetic changes in HSC which affect Runx1-regulated genes. Disrupted MUFA production and reduced metabolic plasticity drives pro-inflammatory macrophage polarisation. These long-term immune effects of HC challenge the notion that further cholesterol lowering will eliminate residual risk of ASCVD, and highlight the potential for identifying rational new therapeutic targets for ASCVD residual risk reduction.

---

**Figure Sup 1:**
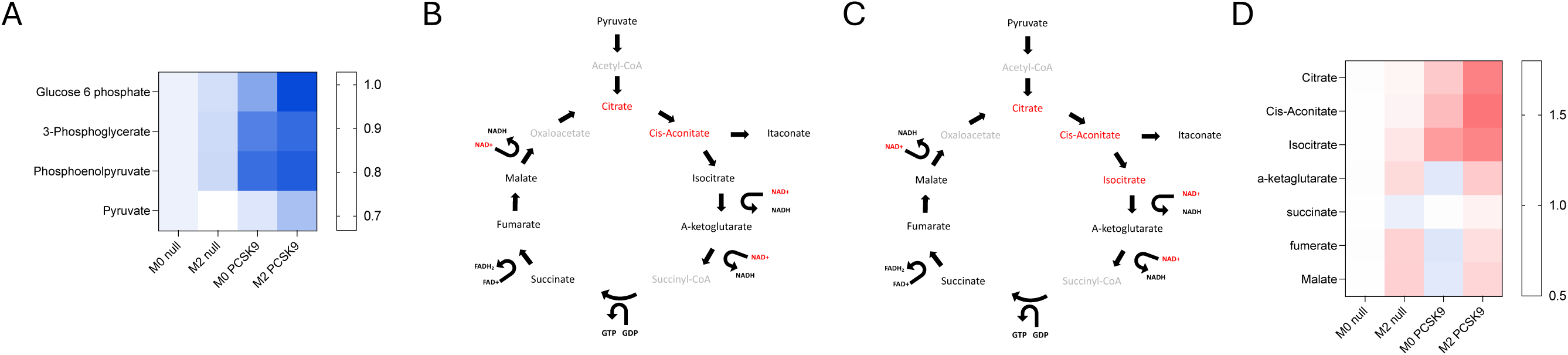
M(IL-4) BMDM from HC mice exhibit reduced central carbon metabolism.

**Figure Sup 2:**
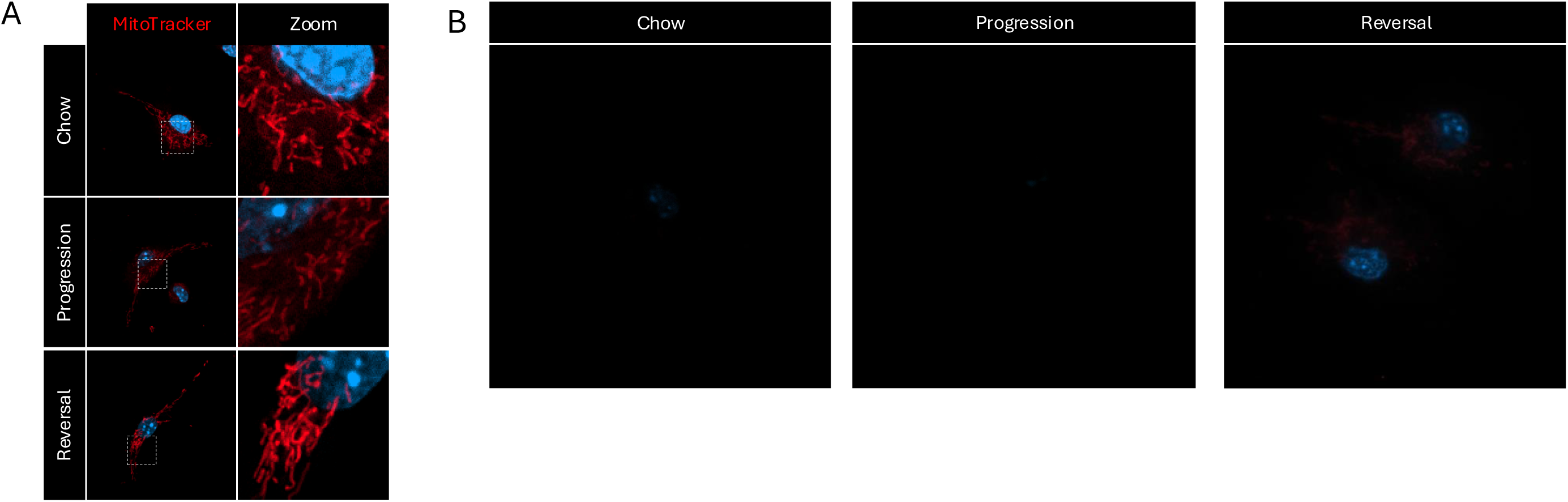
High cholesterol does not alter M(IL-4) BMDM mitochondrial network.

**Figure Sup 3:**
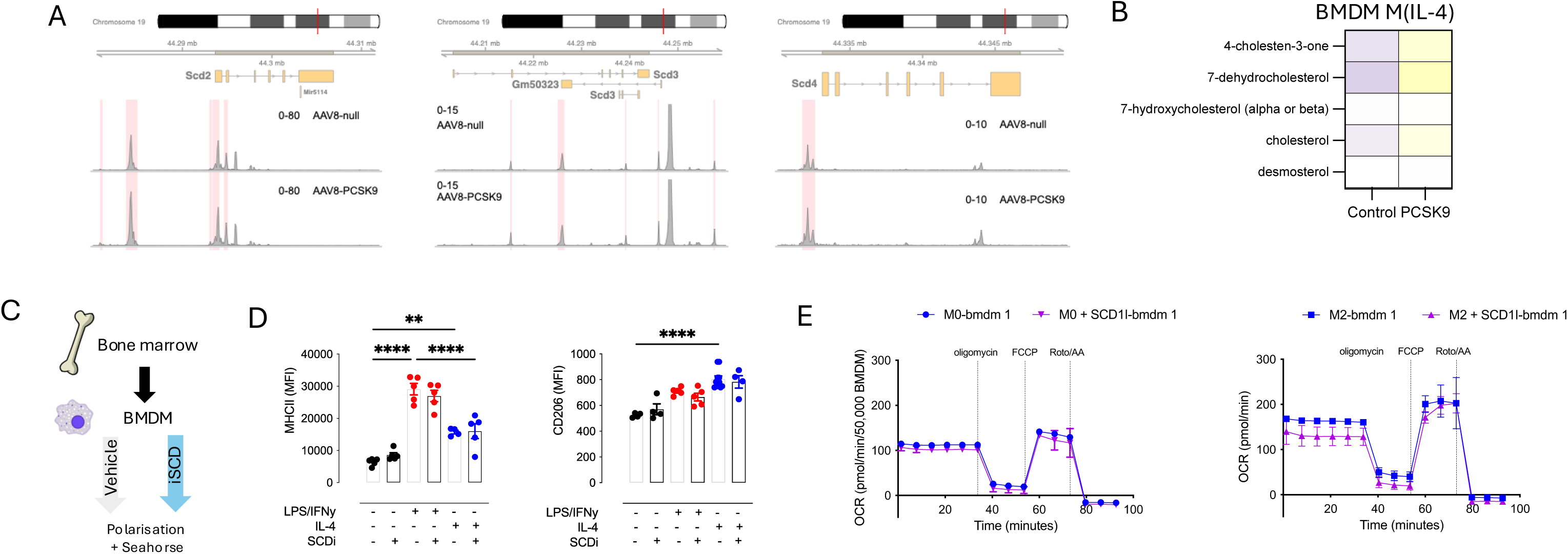
SCD alters monocytes and macrophage MUFA abundance.

**Figure Sup 4:**
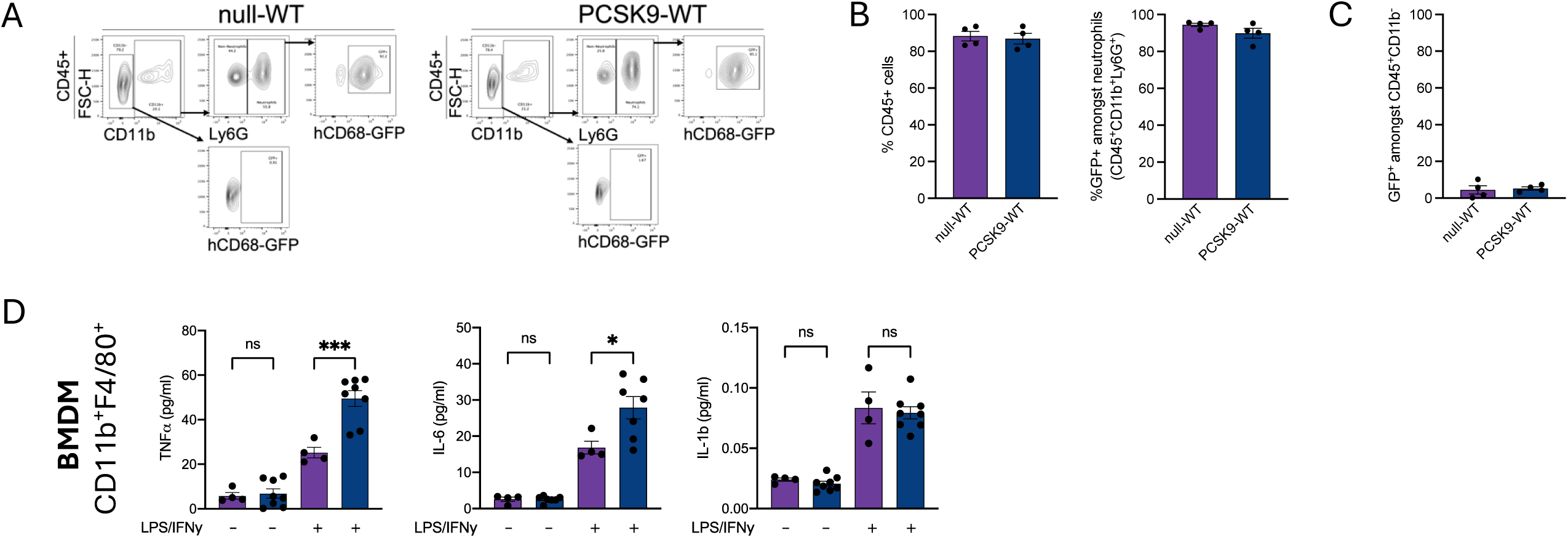
BMDM from chimeric hypercholesterolaemic mice retain an altered macrophage phenotype even after normalization of cholesterol.

**Figure Sup 5:**
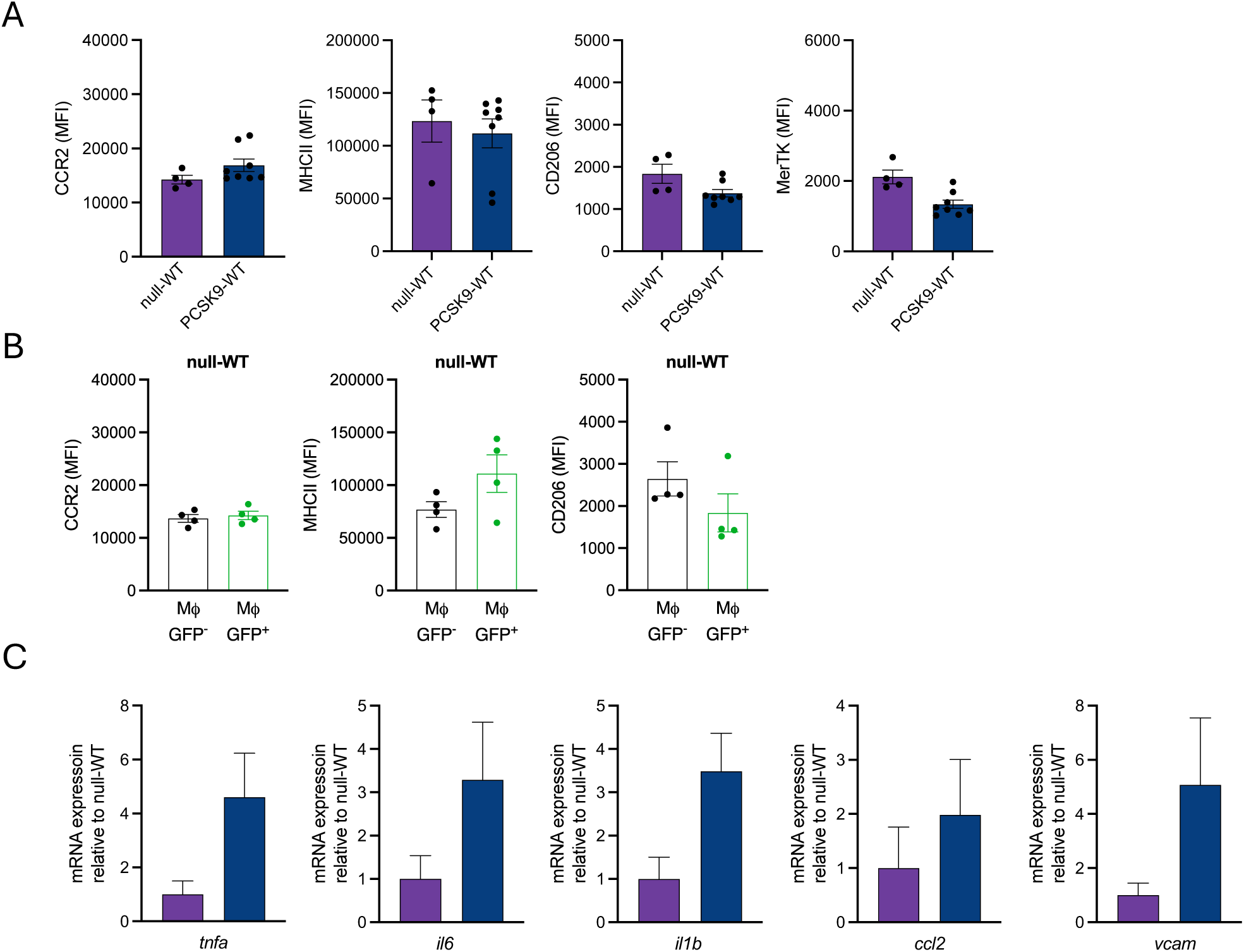
Transplantation of BM from hypercholesterolaemic mice confers a pro-inflammatory phenotype in recipient adipose tissue.

